# TRPV4-mediated Mechanotransduction Regulates the Differentiation of Valvular Interstitial Cells to Myofibroblasts: Implications for Aortic Stenosis

**DOI:** 10.1101/2024.11.05.622116

**Authors:** Pritha Mukherjee, Manisha Mahanty, Bidisha Dutta, Suneha G. Rahaman, Karunakaran R. Sankaran, Shaik O. Rahaman

## Abstract

As aortic valve stenosis (AVS) progresses, the valve tissue also stiffens. This increase in tissue stiffness causes the valvular interstitial cells (VICs) to transform into myofibroblasts in response. VIC-to-myofibroblast differentiation is critically involved in the development of AVS. Herein, we investigated the role of mechanosensitive Ca^2+^-permeant transient receptor potential vanilloid 4 (Trpv4) channels in matrix stiffness- and transforming growth factor β1 (TGFβ1)-induced VIC-myofibroblast activation. We confirmed Trpv4 functionality in primary mouse wild-type VICs compared to Trpv4 null VICs using live Ca^2+^ influx detection during application of its selective agonist and antagonist. Using physiologically relevant hydrogels of varying stiffness that respectively mimic healthy or diseased aortic valve tissue stiffness, we found that genetic ablation of Trpv4 blocked matrix stiffness- and TGFβ1-induced VIC-myofibroblast activation as determined by changes in morphology, alterations of expression of α-smooth muscle actin, and modulations of F-actin generation. Our results showed that N-terminal residues 30-130 in Trpv4 were crucial for cellular force generation and VIC-myofibroblast activation, while deletion of residues 1-30 had no noticeable negative effect on these processes. Collectively, these data suggest a differential regulatory role for Trpv4 in stiffness/TGFβ1-induced VIC-myofibroblast activation. Our data further showed that Trpv4 regulates stiffness/TGFβ1-induced PI3K-AKT activity that is required for VIC-myofibroblast differentiation and cellular force generation, suggesting a mechanism by which Trpv4 activity regulates VIC-myofibroblast activation. Altogether, these data identify a novel role for Trpv4 mechanotransduction in regulating VIC-myofibroblast activation, implicating Trpv4 as a potential therapeutic target to slow and/or reverse AVS development.

## INTRODUCTION

Aortic valve stenosis (AVS) is a progressive disease characterized by fibrosis, inflammation, calcification, and stiffening and thickening of the aortic valve leaflets which leads to disrupted blood flow and eventual left ventricular pressure-overload^1,2^. Symptomatic AVS can lead to heart failure and death within 2 to 5 years if left untreated^3-5^ exemplifying its high mortality rates^3-5^. AVS poses a vexing medical challenge because the only current clinical treatment available is an invasive and risky valve replacement^1,6^. Elucidating the molecular mechanisms underlying the development and progression of AVS is crucial for the development of effective and noninvasive therapeutic strategies to eliminate or reduce AVS.

Valvular interstitial cells (VICs), which are fibroblast-like cells and constitute the primary cellular component of aortic valve leaflets, play a crucial role in maintaining tissue homeostasis, repair, and upkeep^7-9^. Typically, routine mechanical and/or biochemical disruptions can trigger VICs to transform into myofibroblasts. These VIC-myofibroblasts are marked by the formation of stress fibers and exhibit enhanced migration, proliferation, and secretion of extracellular matrix (ECM) components^10^. Under normal conditions, myofibroblasts revert to a quiescent fibroblast state after injury resolution. However, in cases of AVS, myofibroblast activation continues, leading to excessive ECM buildup and remodeling, with subsequent tissue stiffening^10^. Emerging data support a role for stiffness of the extracellular and intracellular matrix in numerous cellular processes including gene expression, inflammation, and cell differentiation^11-21^. VIC-myofibroblast activation and stiffening of the aortic valve are considered primary drivers of AVS development and progression^22-27^. As previously reported by our lab and others, various fibroblast and macrophage activities, including migration and myofibroblast activation, are sensitive to matrix stiffness. This suggests stiffening of valve leaflets may play a regulatory role in AVS development and additionally implies the participation of a cellular stiffness sensor ^11-13,14,20,21,23,28-31^. Published work by our lab also shows that mechanosensitive Trpv4 channels play regulatory roles in fibrosis development in other organs *in vivo* and control fibroblast activation, suggesting that Trpv4 may act as a stiffness sensor in regulating AVS^14,16-18,20,21,31^. Thus, it is essential to verify the identity of the cellular stiffness sensor in VICs and identify the stiffness-induced fibrotic signals that mediate the link between matrix stiffness and AVS development.

Trpv4 is expressed in various cell types including fibroblasts^14,20,21,28,32-37^. Our lab and others have shown that Trpv4 is activated by a range of soluble and mechanical stimuli including substrate stiffness and cytokines^14,16-18,20,21,28,31-37^. In addition to its transmembrane structure which includes a ligand-binding pore, Trpv4 contains several regulatory domains including PI3K recognition domains^32,33^. In mice, Trpv4 deficiency is associated with alterations in pressure responses, osteogenesis, fibrosis, and inflammation^14,20,21,28,32-37^. Trpv4 is also linked to human diseases including skeletal dysplasia and sensory and motor neuropathies^32-34^. Moreover, our data show that Trpv4 deletion inhibited both the substrate stiffness- and growth factor-induced activation of the PI3K-AKT pathway in skin epithelial cells^16,17^. Research on Trpv4’s role in the aortic valve is limited, with studies showing higher Trpv4 protein levels in diseased valves compared to healthy ones, though the specific mechanisms remain unclear^38^. Although aortic valve tissues were not directly studied, increased Trpv4 in the myocardium of diabetic mice is linked to cardiac fibrosis. This alludes to a potentially similar role in AVS, influenced by ECM regulation of Trpv4 in VICs^38,14,17,20,38-40^. Recent research has shown that Trpv4 regulates valve myofibroblast activation and proliferation, suggesting Trpv4 as a candidate mechanosensor in VICs^41^. Nonetheless, the molecular mechanisms by which Trpv4-generated mechanosensing signals (mechanotransduction) are transduced into cells to drive VICs to myofibroblast differentiation remain unknown. In this regard, we found that 1) Trpv4 is functional in mouse VICs; 2) genetic ablation of Trpv4 prevented matrix stiffness and TGFβ1-induced VIC myofibroblast activation; 3) N-terminal residues 30-130 in Trpv4 are critical for cellular force generation and VIC myofibroblast differentiation; and 4) Trpv4 regulates matrix stiffness-induced PI3K-AKT activity to modulate VIC myofibroblast differentiation and cellular force generation, identifying a potential mechanism by which Trpv4 activity regulates VIC-myofibroblast activation. Together, we identify Trpv4 mechanotransduction as a critical regulator of VIC-myofibroblast activation, implicating Trpv4 as a novel target for therapeutic intervention.

## MATERIALS AND METHODS

### Reagents and antibodies

For Western blot and immunofluorescent microscopy analysis the following antibodies were purchased from Cell Signaling Technologies, Danvers, MA: p-Smad2 (3108s), Smad2 (5339s), p-Smad3 (9520s), Smad3 (9523s), p-AKT (9271s), AKT (9272s), p-ERK1/2 (9101s), ERK1/2 (4695s), p-Jnk (4668s), Jnk (9252s), p-P38 (9211s), P38 (9212) and Beta-actin (4970s). Anti-alpha smooth muscle actin antibody (α-SMA) was purchased from Sigma (A2547-2ML) (St. Louis, MO). Anti-Trpv4 antibody (ACC-034) and blocking peptide (BLPCC-034) were purchased from Alomone labs. Pierce RIPA (89900) buffer and Halt protease and phosphatase inhibitor combination (78442) from Thermo Scientific were used to prepare whole cell lysate. Thermo Scientific’s Alexa Fluor 488 conjugated secondary IgGs (A11001) and Alexa fluor phalloidin 594 (A12381) was used for the fluorescent detection. ProLong Gold Diamond 4′,6-diamidino-2-phenylindole (DAPI) mounting media was purchased from Invitrogen (#P36962). Trpv4 antagonist GSK2193874 (GSK219) (cat# SML0942), Trpv4 agonist GSK1016790A (GSK101) (cat# 6433) and PI3K antagonist LY-294002 (LY) (L9908) were purchased from Sigma. TGFβ1 (#240-B) was purchased from R&D Systems. For removing cell adhesion and passaging, Gibco 0.25% Trypsin EDTA solution was used (#25-300-120). Cells were cultured using Gibco’s Dulbecco’s modified Eagle’s medium (DMEM) (11995-011) and heat-inactivated fetal bovine serum (FBS) (10082-147), and antibiotic/antimycotic solution purchased from Sigma (A5955). Culture grade BSA (SH30574.02) was obtained from Hyclone. For traction force microscopy (TFM) studies, Softwell (SW24G-COL-25-STO.2R-ER) TFM plates with embedded fluorescent beads were purchased. Collagen type I (cat#-804622-20ML) and Collagenase-II solution (cat#-C2-22-1G) were purchased from Sigma. Matrigen softslip 24 well culture plates with removable coverslips coated with 1 and 50 kPa stiffness (SS24-COL-1-EA, SS24-COL-50-EA) polyacrylamide (PA) hydrogel was purchased for stiffness-related studies. Adenovirus vector (empty vector), Adeno-RGD-mouseTRPV4 WT, Adeno-GFP-TRPV4, and 3 deletion constructs (Ad-TRPV4-del-1-30, Ad-TRPV4-del-1-130 and Ad-TRPV4-del-100-130) were prepared by Vector Biolabs. For calcium influx studies, FLIPR Calcium 6 assay kit was purchased from Molecular Devices and Calbryte 590 (20700) was purchased from ATT-bioquest.

### Animals

Trpv4 knockout (KO) mice on C57BL/6 background, originally created by Dr. Makato Suzuki (Jichi Medical University, Tochigi, Japan), was obtained from Dr. David X. Zhang (Medical College of Wisconsin, Milwaukee, WI). The congenic wild type (WT) mice were purchased from Charles River Labs. All mice were housed and bred in a temperature and humidity-regulated, germ-free environment with food and water *ad libitum*. For all the experiments, Institutional Animal Care and Use Committee (IACUC) guidelines were followed and animal protocols were approved by the University of Maryland College Park review committee.

### Valvular interstitial cell culture and maintenance

We have harvested VICs from mice hearts following the protocol from JOVE Journal^42^. VICs were extracted from WT and Trpv4 KO mice after dissecting the heart and valve areas under a light microscope. Extracted valve leaflets were treated with collagenase II and incubated for 1 h at 37°C. Valvular endothelial cells were removed by vortexing leaflets vigorously at maximum speed for 60 seconds. After the incubation the tissues were minced using a sterile scalpel and incubated in collagenase II for additional 15 minutes. After the second incubation the tissues and cell mixture were pipetted several times to loosen the cells from the tissue surface following a wash with sterile PBS and resuspended in complete 10% FBS containing DMEM (supplemented with antibiotic and antimycotic solution) on collagen-coated culture plates. Cells were grown with fresh DMEM medium every 48 h. We used 0.25% trypsin EDTA solution for passaging the cells, and 10% DMSO plus 90% FBS solution was used to freeze the cells in liquid nitrogen until further usage. All the experiments are performed using cells within passage number 3-8. Cells were maintained in 10% FBS containing DMEM for all the experiments related to adenovirus transfection along with TGFβ1 treatment. For TGFβ1 treatment without any adenovirus, we used 1% BSA containing DMEM.

### Traction force microscopy (TFM)

VICs were plated on collagen-coated (10 μg/ml) 25 kPa PA hydrogels containing 0.2 μm fluorescent beads in 24-well plates. WT and Trpv4 KO VICs were allowed to grow overnight at 37°C. Subsequently, the cells were subjected to treatment with TGFβ1 (5 ng/ml) for 48 hours. For experiments with adenovirus, Trpv4 KO VICs were transfected by adenovirus constructs with or without Trpv4 antagonist GSK219 or PI3K inhibitor LY treatment. Using a microscope with a 20x objective, fluorescent images of beads were captured both before and after lysing the cells using a 0.6% SDS solution. Bright field picture of the cells was taken after capturing the initial state of the fluorescent beads. To quantify the displacement and cellular traction force generated, we utilized the TractionForAll® software^43^.

### Calcium influx study

Fluorescence imaging of VICs was done using Perkin Elmer spinning disc laser confocal microscope fitted with 20x objective lens. WT and Trpv4 KO VICs were cultured on 35 mm glass bottom plates with Ad-Trpv4 or Ad-vec for 48 hours. For the Ca^2+^ influx measurement, the cells were washed with HBBS buffer (Corning, cat# 21023105) twice and incubated with Calbryte 590 AM dye in HBBS for 1 h at 37°C. Following two more washes, transient fluorescence measurement was performed by perfusing the cells with buffer containing 100 nM GSK 1016790A (GSK101), a known Trpv4 agonist, and fluorescence was measured for 8 mins. The fluorescence intensity was analyzed and quantified using Image J. In the second method of Ca^2+^ influx study, we used FlexStation 3 instrument to measure GSK101-induced Trpv4-induced Ca^2+^ influx in WT and Trpv4 KO VICs cultured in 96-well black wall clear bottom plate, with or without TGFβ1(5 ng/ml) treatment for 48 hours. Prior to the experiment cells were pre-treated with or without GSK219 for 1 hour. FLIPR calcium 6 Assay Kit was used to quantify changes in the intercellular Ca^2+^ influx. VICs were washed and incubated for 45 min with FLIPR kit reagent (Calcium 6 dye in 1x HBSS containing 20 mM HEPES and 2.5 mM probenecid) at 37°C and then transferred to the FlexStation 3 instrument for measuring the Ca^2+^ influx^14, 20^.

### Western blot analysis

Whole cell extracts were prepared from WT and Trpv4 KO VICs cultured on plastic tissue culture plates coated with collagen, with or without TGFβ1 treatment for 48 h using RIPA buffer containing protease and phosphatase inhibitors cocktail. Cell lysates were separated on a 10% SDS-PAGE, transferred to a PVDF membrane, and probed with antibodies against p-Smad2 (1:1000), Smad2 (1:1000), p-Smad3 (1:1000), Smad3 (1:1000), α-SMA (1:1000), p-AKT (1:1000), AKT (1:1000), p-ERK (1:1000), ERK (1:1000), p-P38 (1:1000), P38,(1:1000), p-JNK (1:1000), JNK (1:1000) or Actin (1:3000), respectively. The blots were visualized using anti-rabbit (1:2000) or anti-mouse (1:2000) horseradish peroxidase-conjugated secondary IgG and developed in UVP Biospectrum. Images were quantified using UVP data processing software.

### Overexpression of full length and mutant Trpv4 gene in Trpv4 KO VICs by adenovirus construct transfection

To conduct gain-of-function studies in Trpv4 KO VICs, cells were transfected with full-length WT Ad-Trpv4 and three deletion constructs of different lengths of mouse Trpv4 gene (Ad-Trpv4-Δ1-30, Ad-Trpv4-Δ1-130, and Ad-Trpv4-Δ100-130). To verify the success of transfection efficiency of adenovirus constructs in the VICs, GFP-tagged Ad-Trpv4 was used at a concentration of 10^7^ plaque forming unit (PFU). Trpv4 KO VICs were seeded on cell culture plates or TFM plates with DMEM, 10% FBS, and transfected with adenovirus Trpv4 constructs or empty vector control (10^7^ plaque-forming units/ml) for 24 h followed by a fresh medium replacement and another 24-48 h of incubation with complete DMEM with or without TGFβ1 (5 ng/ml) treatments.

### Immunofluorescence microscopy analysis

To analyze the production of α-SMA and F-actin, WT and Trpv4 KO VICs were cultured on collagen-coated glass coverslips with or without the TGFβ1 (5 ng/ml) treatment for 48 hours. To determine the effect of matrix stiffness on the production of α-SMA and F-actin, VICs were cultured on 1 and 50 kPa PA hydrogel-coated coverslips with or without TGFβ1 treatment for 48 hours. Cells were fixed with 4% paraformaldehyde and were incubated with 3% FBS plus 0.1% triton-x containing blocking solution. Anti-α-SMA antibody was used in a 1:100 ratio and the secondary antibody was used in a 1:200 Ratio. To label F-actin, Alexa-Fluor-labeled phalloidin 594 was used. DAPI-containing mounting medium was used to stain the nucleus. To confirm the specificity of the Trpv4 antibody in VICs, WT VICs were cultured on glass coverslips, fixed with 4% paraformaldehyde, and stained with anti-Trpv4 (1:200) primary and anti-Rabbit (1:300) Alexa-fluor conjugated secondary IgG, with or without the presence of Trpv4 blocking peptide.

## RESULTS

### Primary mouse VIC express functional Trpv4 channels that can be activated and inhibited by its selective agonist and antagonist

Published work by our lab and others showed that Trpv4 plays critical roles in fibrosis development in other organs and regulate fibroblast activation, indicating that Trpv4 may be involved in regulating VIC-to-myofibroblast activation as related to AVS^14,16,20,40,41,44^. We hypothesized that mouse primary VICs express functional Trpv4 channels, and its channel function could be modulated by selective agonists and antagonists. First, using immunofluorescence microscopy, we confirmed that Trpv4 protein is present in VICs cultured on tissue culture plates (Fig. 1A). Blocking of Trpv4-specifc signals by a Trpv4-targeted peptide confirmed the specificity of the Trpv4 antibody (Fig. 1A). Next, we used real-time Ca^2+^ influx imaging to test the functionality of Trpv4 channel and detect cytoplasmic Ca^2+^ levels induced by Trpv4 agonist GSK101 in VICs. Ca^2+^ influx levels were increased 3-fold in VICs stimulated by GSK101 compared to unstimulated cells (Fig. 1B). Interestingly, further increased levels of Ca^2+^ influx was detected in TGFβ1-stimulated VICs compared to no TGFβ1control, suggesting a critical role of Trpv4 in TGFβ1-induced Ca^2+^ influx in VICs. Additional results showed that TGFβ1-dependent Ca^2+^ influx was completely blocked by treatment with a Trpv4 antagonist GSK219, suggesting that TGFβ1-elicited Ca^2+^ influx was mediated by Trpv4 (Fig. 1B).

**Figure 1.**
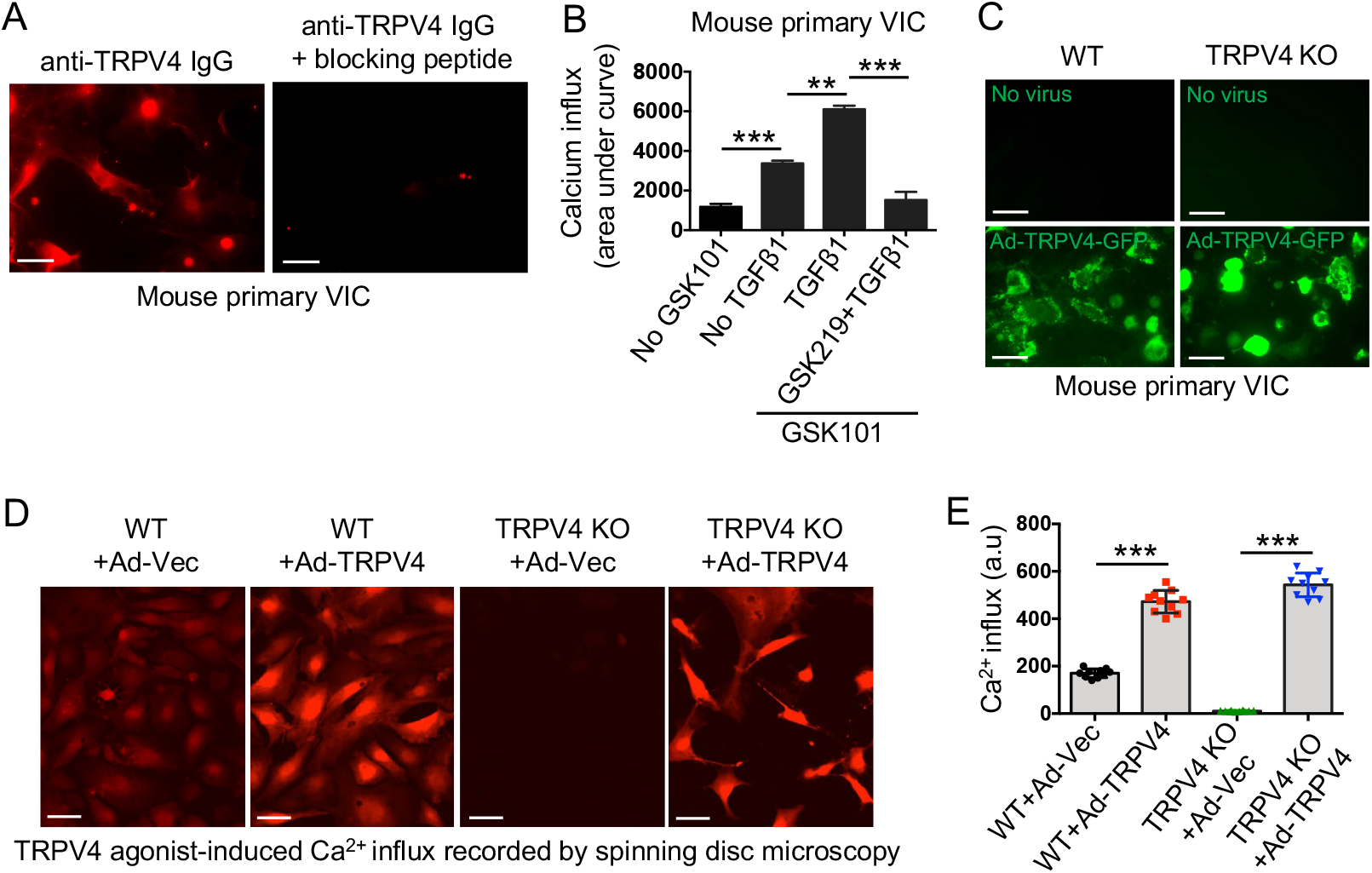
Primary mouse VIC express functional Trpv4 channels/receptors. **A**. VICs from WT mouse was immunostained with Trpv4 antibody (red). Trpv4 antibody pre-incubated with Trpv4 blocking peptide (right panel) was used to show specificity of Trpv4 IgG. **B**. VICs (∼20,000 cells per well) were seeded on collagen-coated (10 μg/ml) 96-well plastic plate. Ca^2+^ influx is shown by are under curve measuring ΔF/F (Max-Min). FlexStation 3 recording of Calcium 6 dye-loaded WT VIC monolayers with or without TGFβ1 treatment show effect of Trpv4 agonist GSK101 or antagonist GSK219 on Ca^2+^ influx. \*\**p* < 0.01; \*\*\**p* < 0.001; Student’s t-test. **C**. Fluorescence microscopy images show overexpression of Trpv4 proteins in WT and Trpv4 KO VICs transfected with GFP tagged Ad-Trpv4 for 48 hours. **D**. Spinning disc confocal microscopy images show recording of Ca^2+^ influx (red signals) in GSK101-exposed WT and Trpv4 KO VICs transfected with control Ad-Vec or Ad-Trpv4 constructs. **E**. Bar graph show quantitation of results (mean ± SEM) from Figure 1D. n = 50 cells per condition. \*\*\**p* < 0.001; Student’s t-test.

We confirmed successful overexpression of WT Trpv4 (Ad-TRPV4-GFP) in adenovirus construct-transfected WT and Trpv4 KO VICs through fluorescence microscopy showing overexpression of Trpv4 proteins specifically in membranes (Fig. 1C). To determine the functionality of the adenovirus construct-derived Trpv4 channel in WT and Trpv4 KO VICs, we assayed Ca^2+^ influx induced by GSK101 in VICs transfected with Ad-Trpv4, or control vector constructs. Ca^2+^ influx was increased to approximately 2.5-fold in Ad-Trpv4 transfected VICs compared to vector control cells (Figs. 1D & E). Altogether, our findings suggested that both mouse primary VICs and VICs transfected with Ad-Trpv4 express functional Trpv4 channels which can be activated and inhibited by Trpv4-selective agonist and antagonist.

### Trpv4 regulates both TGFβ1- and matrix stiffness-induced VIC-to-myofibroblast differentiation

Published work from our lab and others suggest a link between Trpv4 activation and TGFβ1 signals that mediate myofibroblast differentiation^14,20,40,41,44^. To test our hypothesis that Trpv4 activation might regulate VIC-to-myofibroblast differentiation, VICs from WT and Trpv4 KO mice were seeded on collagen-coated plastic plates and treated with TGFβ1. They were then compared for acquisition of myofibroblast-related biochemical changes. We found that compared to untreated WT controls, TGFβ1-pretreated WT VICs showed upregulated F-actin generation along with upregulated α-SMA expression (Figs. 2A-C), confirming the profibrotic effect of TGFβ1. In contrast, results of immunofluorescence analysis indicated that Trpv4 KO VICs treated or not with TGFβ1 showed significant reductions in expression of α-SMA and generation of F-actin signals compared to TGFβ1-treated WT VICs (Figs. 2A-C). These results suggest that Trpv4 is required for TGFβ1-induced VIC-to-myofibroblast differentiation.

**Figure 2.**
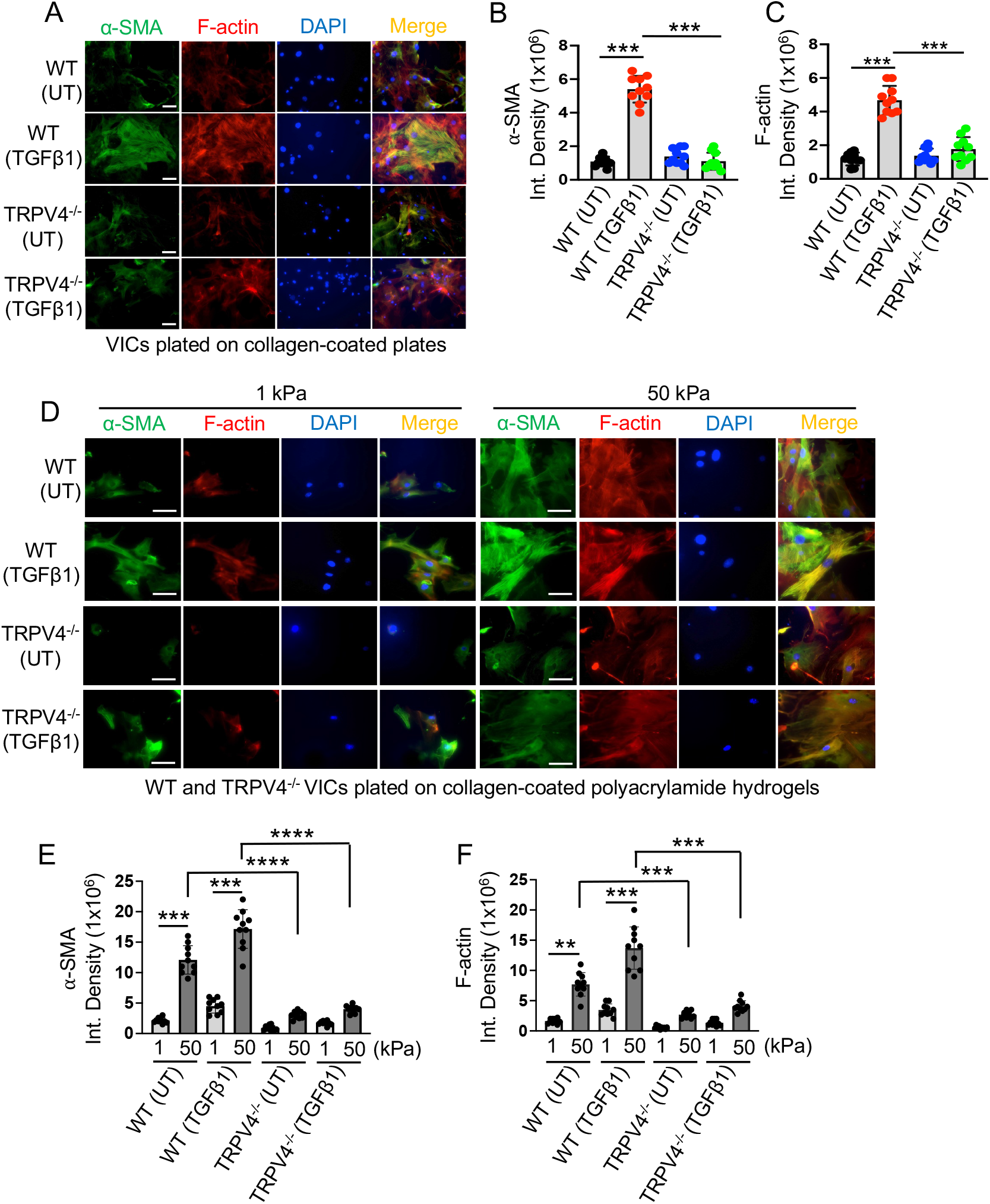
Trpv4 is required for both TGFβ1- and matrix stiffness-induced myofibroblast differentiation in VICs. **A**. WT and Trpv4 KO VICs were plated on collagen-coated plastic plates and incubated with or without TGFβ1 (5 ng/ml) for 48 h. Representative images from five different fields per condition shows morphological changes and co-localization of ɑ-SMA (green) and F-actin (red) with DAPI (blue) staining in VICs. **B** & **C**. Quantitation of results from Figure 2A. Data are expressed as mean ± SEM, n = 20 cells/condition; UT = untreated; \*\*\**p* < 0.001, 1-way ANOVA. **D**. WT and Trpv4 KO VICs were plated on collagen-coated soft (1 kPa) and stiff (50 kPa) polyacrylamide hydrogels and incubated with or without TGFβ1 (5 ng/ml, 48 h). Representative images from five different fields per condition showing morphological changes and co-localization of ɑ-SMA (green) and F-actin (red) with DAPI (blue) staining in WT and Trpv4 KO VICs in response to increasing matrix stiffness and TGFβ1 treatment. **E** & **F**. Quantitation of results from Figure 2D. Data are expressed as mean ± SEM, n = 20 cells/condition; UT = untreated; \*\**p* < 0.01; \*\*\**p* < 0.001, \*\*\*\**p* < 0.0001, 1-way ANOVA.

Previous reports from our lab and others showed that increases in matrix stiffness drive TGFβ1-mediated myofibroblast differentiation in other cell types^14,20,40,41,44^. To ascertain the role of Trpv4 in VIC-to-myofibroblast differentiation as a result of increasing matrix stiffness alone or in combination with TGFβ1, we seeded VICs on soft (1 kPa) or stiff (50 kPa) collagen-coated polyacrylamide (PA) hydrogels treated with or without TGFβ1. We then assessed the occurrence of myofibroblast differentiation in Trpv4 KO and WT VICs. Immunofluorescence analysis showed that TGFβ1 was unable to drive myofibroblast differentiation in WT VICs under soft conditions (normal tissue stiffness) (Figs. 2D-F). In contrast, under stiff matrix conditions (fibrotic tissue stiffness) without TGFβ1 treatment, WT VICs differentiated to myofibroblasts as confirmed by increased production of α-SMA, F-actin, and stress fibers (Fig. 2D). Intriguingly, the addition of TGFβ1 further augmented the matrix stiffness-induced myofibroblast generation in WT VICs under stiff matrix conditions (Figs. 2D-F). However, stiff matrix and TGFβ1 did not induce myofibroblast differentiation in Trpv4 KO VICs (Figs. 2D-F). These results indicate that Trpv4 mechanosensing activity is required for both matrix stiffness and TGFβ1-induced VIC-to-myofibroblast differentiation.

### N-terminal residues 30 to 130 in Trpv4 is critical in regulation of TGFβ1-induced VIC-to-myofibroblast differentiation

The N-terminal regions of Trpv4, recognized for their role in channel activation through biomechanical stimuli like osmolarity and stiffness, were examined for their potential to regulate VIC-to-myofibroblast differentiation in response to TGFβ1^20,45^. For this, three Trpv4 mutants with progressive N-terminal deletions (Ad-Trpv4-Δ1-30, Ad-Trpv4-Δ1-130, and Ad-Trpv4-Δ100-130) were produced using an adenovirus expression system (Fig. 3A)^20^. These mutant constructs, along with the full-length-WT Trpv4 (Ad-Trpv4) and an empty vector (Ad-Vec), were introduced into Trpv4 KO VICs. Post-TGFβ1 treatment, VICs expressing the WT Trpv4 or Ad-Trpv4-Δ1-30 construct showed increased VIC myofibroblast differentiation compared to the empty vector control. Notably, Trpv4 overexpression-dependent TGFβ1-induced myofibroblast differentiation in Trpv4 KO VICs was completely blocked by Trpv4 antagonist GSK219. These results suggest a direct role of Trpv4 in this process (Figs. 3B-D). The high level of VIC-to-myofibroblast differentiation was suppressed in cells transfected with Ad-Trpv4-Δ1-130 or Ad-Trpv4-Δ100-130 indicating the critical role of the Trpv4 30-130 region in regulating TGFβ1-induced VIC myofibroblast differentiation (Figs. 3B-D). All together, these results suggested that residues 30-130 in Trpv4 were crucial for VIC myofibroblast differentiation, while truncation of residues 1-30 had no noticeable negative effect on myofibroblast differentiation.

**Figure 3.**
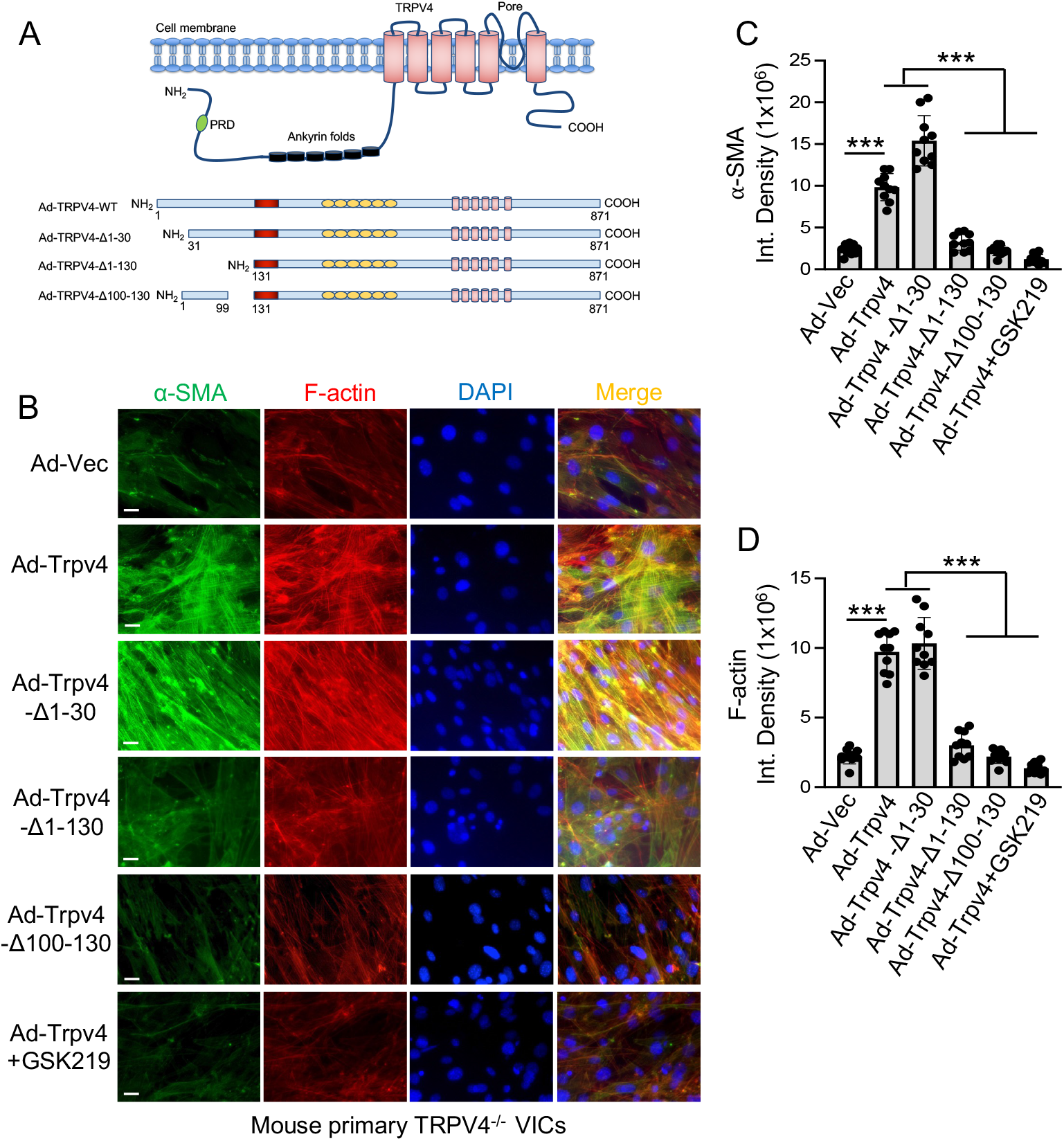
Differential requirement of Trpv4 domains for TGFβ1-induced VIC-to-myofibroblast differentiation. **A**. Schematic diagram of full-length Ad-TRPV4-WT and three Trpv4 constructs with deletions in N-terminal residues (Adeno-TRPV4-Δ-1-30, Adeno-TRPV4-Δ-1-130 and Adeno-TRPV4-Δ-100-130). PRD: proline rich domain. **B**. Representative images from five different fields per condition showing morphological changes and co-localization of ɑ-SMA (green) and F-actin (red) with DAPI (blue) staining in TGFβ1 (5 ng/ml)-stimulated Trpv4 KO VICs transfected with Ad-TRPV4-WT (with or without GSK219), Ad-TRPV4-Δ1-30, Ad-TRPV4-Δ1-130, Ad-TRPV4-Δ100-130, or Ad-Vec. **C** & **D**. Quantitation of results from Figure 3B. Data are expressed as mean ± SEM, n = 20 cells/condition; \*\*\**p* < 0.001, 1-way ANOVA.

### Differential requirement of Trpv4 domains for TGFβ1-induced traction force generation in VICs

Cytoskeletal remodeling is essential to various cellular processes, including cell differentiation. The generation of cellular traction force is primarily dependent on F-actin production, a key component of cytoskeletal remodeling^46,47^. Our findings revealed a notable increase in Trpv4-dependent F-actin production on stiffer PA gels compared to softer ones in VIC-myofibroblasts (Figs. 2D-F). To determine if Trpv4-dependent VIC-myofibroblast formation was associated with cellular traction force generation, we employed traction force microscopy to measure force generation in VICs induced by TGFβ1. We found that TGFβ1 stimulation significantly increased force generation in WT VICs compared to Trpv4 KO VICs (Figs. 4A & B), highlighting Trpv4’s essential role in cellular force generation. Collectively, these results suggest that F-actin-dependent cytoskeletal remodeling, and the subsequent increase in cellular force, may be instrumental in Trpv4-dependent VIC-myofibroblast generation.

**Figure 4.**
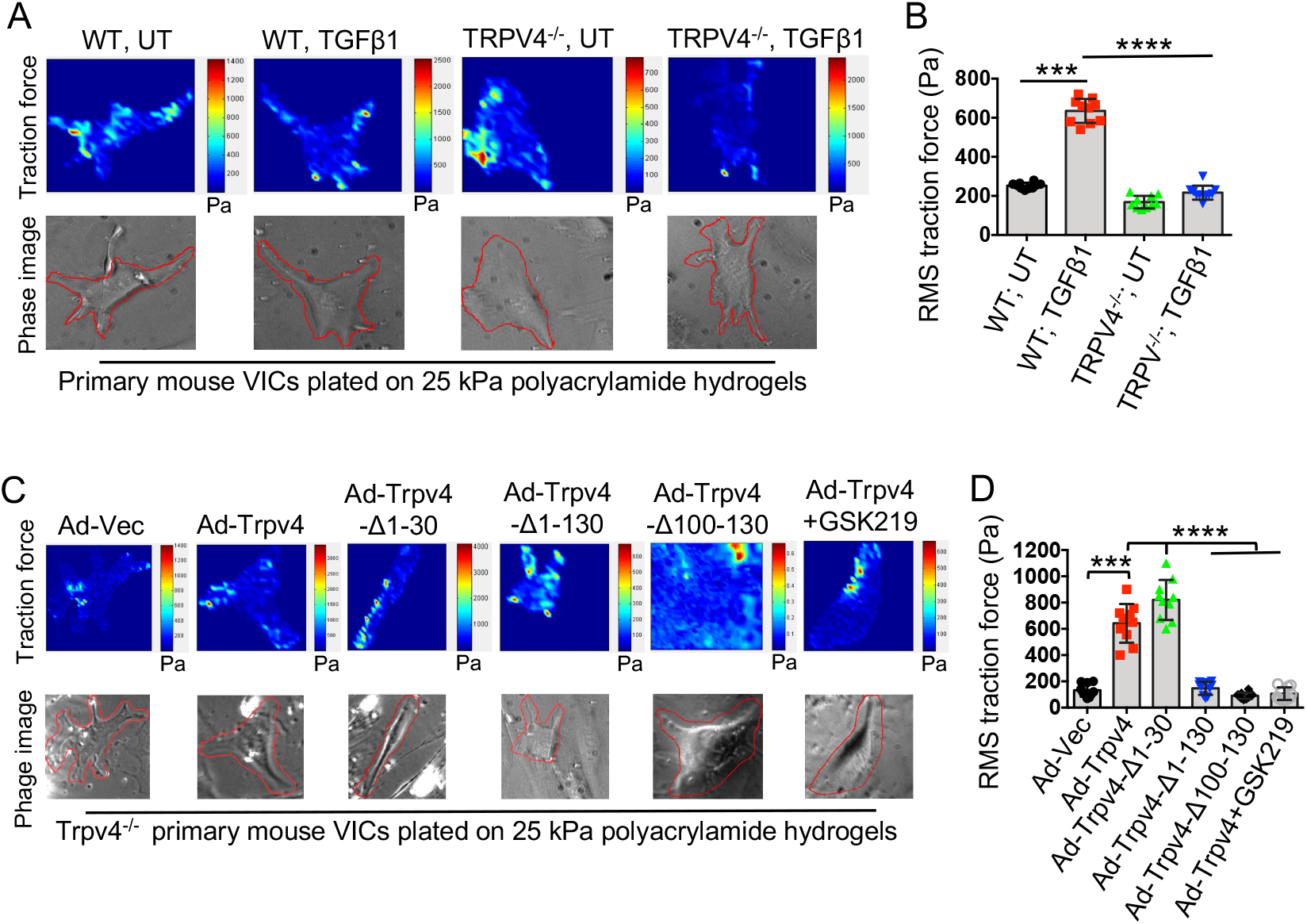
N-terminal residues 30 to 130 in Trpv4 is critical in regulation of TGFβ1-induced traction force generation in VICs. **A**. WT and Trpv4 KO VICs were plated on collagen-coated (10 µg/ml) 25 kPa PA hydrogels containing 0.2 μm fluorescent beads in 24-well plates with or without 5 ng/ml of TGFβ1 for 24 h. Color-coded traction vector maps indicate the magnitude of the traction vector and corresponding phase images show the cells. **B**. Quantitation of results from traction force microscopy assay as shown in Figure 4A. Data are expressed as mean ± SEM, n = 10 cells/condition; \*\*\**p* < 0.001, \*\*\*\**p* < 0.0001, Student’s t-test; RMS: root mean square. **C**. Trpv4 KO VICs transfected with Ad-TRPV4-WT (with or without GSK219), Ad-TRPV4-Δ1-30, Ad-TRPV4-Δ1-130, Ad-TRPV4-Δ100-130, or Ad-Vec were seeded on collagen-coated (10 µg/ml) 25 kPa PA hydrogels containing 0.2 μm fluorescent beads in 24-well plates and were stimulated by 5 ng/ml of TGFβ1 for 24 h. **D**. Bar graph shows quantitation of results from Figure 4C. Data are expressed as mean ± SEM, n = 10 cells/condition; \*\*\**p* < 0.001, \*\*\*\**p* < 0.0001, Student’s t-test; UT: untreated.

We generated gain-of-function by overexpression of Ad-Trpv4-WT, Ad-Trpv4-Δ1-30, Ad-Trpv4-Δ1-130, and Ad-Trpv4-Δ100-130 in Trpv4 KO VICs through transfection of these adenovirus expression constructs. In these experiments, an empty adenovirus vector (Ad-Vec) was used as a negative control. Cells expressing the Ad-Trpv4-WT or Ad-Trpv4-Δ1-30 construct showed increased force generation compared to the empty vector control. Notably, Trpv4 overexpression-dependent force generation in Trpv4 KO VICs was completely blocked by Trpv4 antagonist GSK219. Together, these results suggest a direct role of Trpv4 in cellular force generation (Figs. 4C & D). The degree of force generation was significantly suppressed in cells transfected with the deletion mutants, Ad-Trpv4-Δ1-130 or Ad-Trpv4-Δ100-130, indicating the critical role of the Trpv4 30-130 region in regulating TGFβ1-induced force generation (Figs. 4C & D). All together, these results suggested that residues 30-130 were crucial for force generation in VICs, while truncation of residues 1-30 had no noticeable negative effect on this process.

### Trpv4 channel regulates TGFβ1-induced AKT activation in VICs

Activation of the PI3K/AKT pathway is a central promoter of cell differentiation in various cell types^16-18,48,49^. The PI3K/AKT pathway can be activated in either a TGFβ1-dependent or - independent manner^50-52^. We did not find any increase in TGFβ1-induced phosphorylation of Smad-2/3 or MAPK proteins (Erk, p38, and Jnk) in both WT and Trpv4 KO VICs (Figs. 5A). However, genetic deletion of Trpv4 significantly suppressed phosphorylation of AKT (at serine 473) with or without TGFβ1 stimulation and inhibited expression of α-SMA by TGFβ1, as compared to WT VICs (Figs. 5A-C). This suggests that both basal and TGFβ1-induced activation of AKT is dependent on Trpv4 in VICs. Notably, Trpv4 overexpression-dependent phosphorylation of AKT and expression of α-SMA in response to TGFβ1 in Trpv4 KO VICs was significantly blocked by Trpv4 antagonist GSK219. Together, these results suggest a direct role of Trpv4 in TGFβ1-induced phosphorylation of AKT in VICs (Figs. 4D-F).

**Figure 5.**
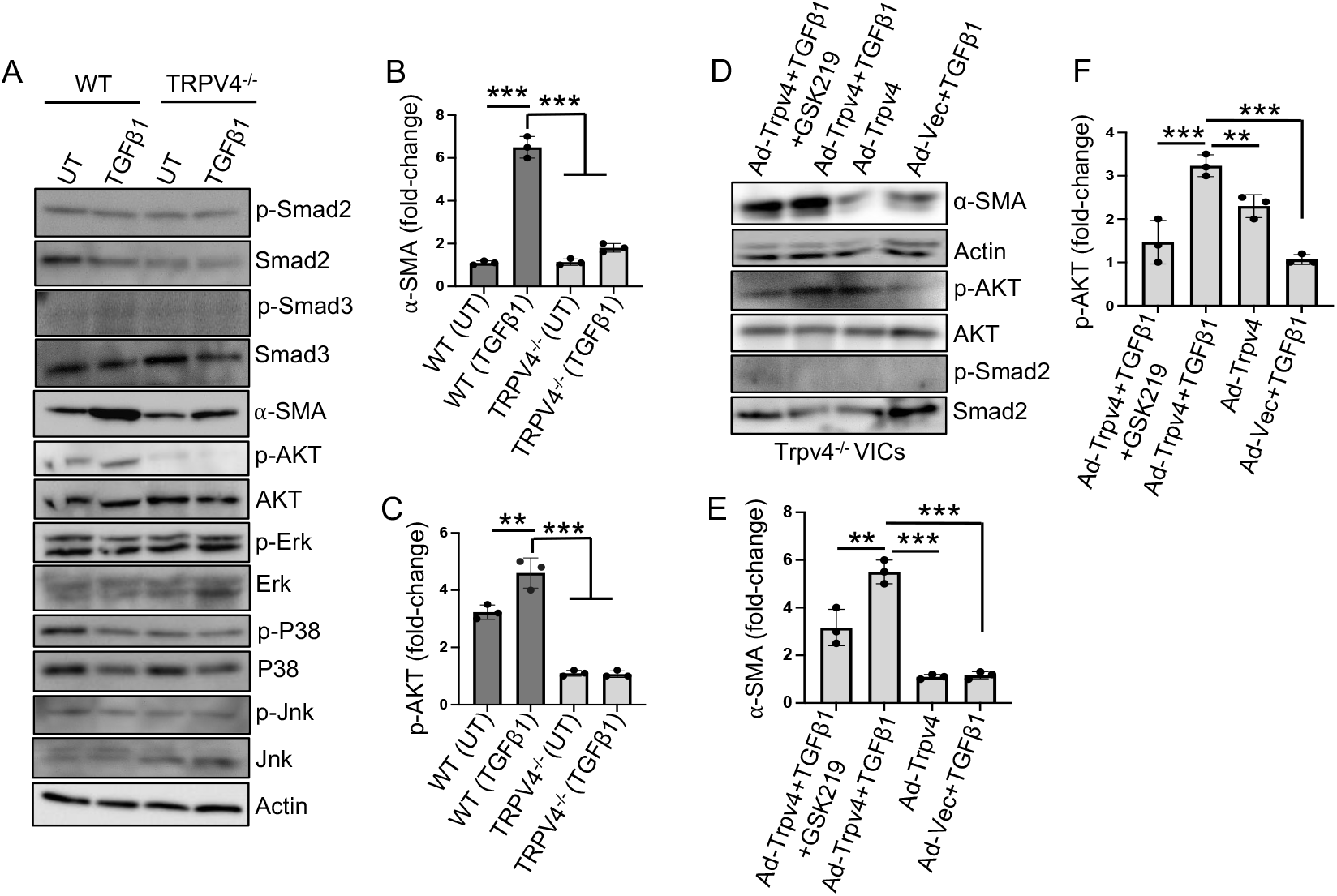
Deficiency of Trpv4 function blocks TGFβ1-induced phosphorylation of AKT in VICs. **A**. VICs were plated on plastic plates coated with collagen (10 µg/ml). Immunoblots show the expression levels of p-Smad2, total Smad2, p-Smad3, total Smad3, *α*-SMA, p-AKT, total AKT, p-Erk, total Erk, p-P38, total-P38. p-Jnk, total Jnk, and actin 48 h after treatment of WT and Trpv4 KO VICs with or without TGFβ1 (5 ng/ml) stimulation. All experiments were repeated at least three times. Graph shows quantification of results from Figure A for *α*-SMA (**B**) and p-AKT (**C**). \*\**p* < 0.01, \*\*\**p* < 0.001, Student’s t-test. **D**. Immunoblots show the expression level of p-Smad2, total Smad2, *α*-SMA, p-AKT, total AKT, and actin 48 h after treatment of Trpv4 KO VICs transfected with Ad-TRPV4-WT (with or without GSK219) or control Ad-Vec in the presence or absence of TGFβ1 (5 ng/ml). All experiments were repeated three times. Graph shows quantification of results from Figure D for *α*-SMA (**E**) and p-AKT (**F**). \*\**p* < 0.01, \*\*\**p* < 0.001.

### Trpv4 regulates matrix stiffness-induced phosphorylation of AKT which is required for VIC- to-myofibroblast differentiation and traction force generation

It has been previously found that activation of the PI3K-AKT pathway is upregulated by stiff matrices in other cell types^16-18,53,54^. We compared phosphorylated AKT (p-AKT) levels between WT and Trpv4 KO VICs grown on soft (1 kPa) and stiff (50 kPa) PA hydrogels. Immunofluorescence analysis showed that stiff matrices caused increased p-AKT levels in WT VICs compared to WT VICs grown on soft matrix (Figs. 6A & B). Intriguingly, the addition of TGFβ1 further augmented the level of p-AKT in WT VICs under stiff matrix conditions (Figs. 6A & B). However, stiff matrix and TGFβ1 did not induce the level of p-AKT in Trpv4 KO VICs (Figs. 6A & B). These results indicate that Trpv4 mechanosensing activity is required for both matrix stiffness and TGFβ1-induced phosphorylation of AKT in VICs. Notably, Trpv4 overexpression-dependent increase in α-SMA and p-AKT levels in Trpv4 KO VICs plated on stiff PA hydrogels were completely blocked by PI3K inhibitor LY (Figs. 6C-E), suggesting a direct role of Trpv4 in VIC myofibroblast differentiation and phosphorylation of AKT under stiff environment. Moreover, we found that Trpv4 overexpression-dependent increase in traction force generation in Trpv4 KO VICs plated on stiff PA hydrogels were significantly blocked by LY (Figs. 6F & G), suggesting a direct role of Trpv4-PI3K axis in cellular force generation under stiff environment.

**Figure 6.**
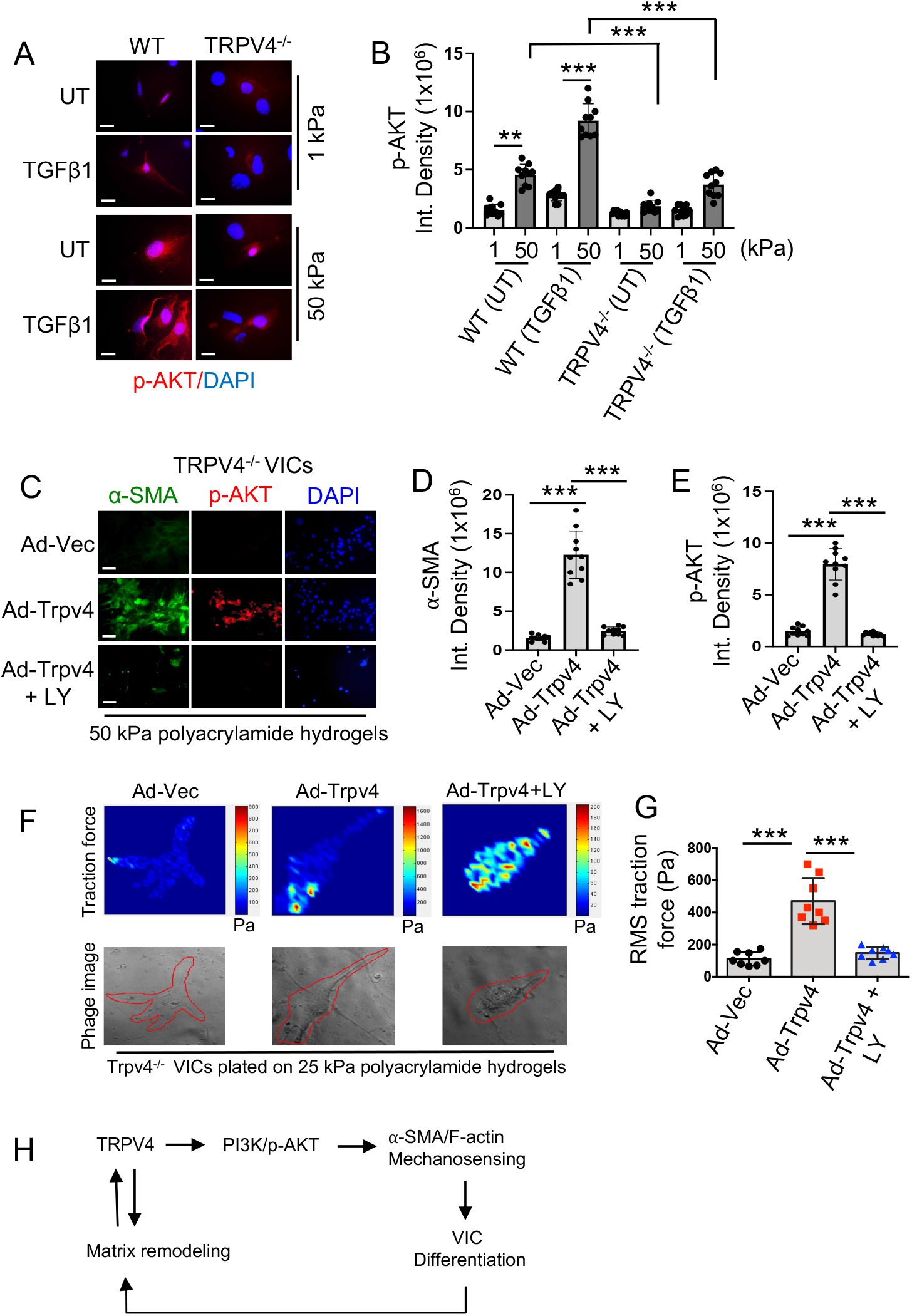
Trpv4-dependent phosphorylation of AKT by matrix stiffening is required for VIC- to-myofibroblast differentiation and traction force generation. **A**. Immunofluorescent microscopy images show the level of p-AKT (red) in WT and Trpv4 KO VICs plated on soft (1 kPa) and stiff (50 kPa) PA hydrogels with or without TGFβ1 (5 ng/ml) treatment for 24h. **B**. Bar graph shows quantification of results from Figure 6A for p-AKT. n = 20 cells/condition; UT = untreated; \*\**p* < 0.01, \*\*\**p* < 0.001, Student’s t-test. **C**. Immunofluorescent microscopy images show the level of p-AKT (red) and ɑ-SMA (green) in Trpv4 KO VICs transfected with Ad-TRPV4-WT (with or without LY) or control Ad-Vec and grown in the presence of TGFβ1 (5 ng/ml) for 24h on stiff (50 kPa) PA hydrogels. **D** & **E**. Bar graph shows quantification of results from Figure 6C for ɑ-SMA (**D**) and p-AKT (**E**). n = 20 cells/condition; \*\*\**p* < 0.001, Student’s t-test. **F**. Color-coded traction vector maps indicate the magnitude of the traction vector and corresponding phase images show the cells. **G**. Quantitation of results from traction force microscopy analysis. Data are expressed as mean ± SEM, n = 10 cells/condition, \*\*\**p* < 0.001, Student’s t-test. **H**. Proposed schematic model showing mechanistic pathway by which TRPV4-dependent mechanotransduction integrates matrix stiffness and soluble signals to induce VIC-to-myofibroblast differentiation via PI3K-AKT pathway.

## DISCUSSION

Key features of AVS include development of fibrosis, inflammation, calcification, and hardening and thickening of the aortic valve leaflets^1,2^. These pathological changes can lead to disrupted blood flow, left ventricular pressure-overload, and even death of the patient^1,2^. VICs, the main cellular component of aortic valve leaflets, play a critical role in maintaining valve homeostasis, repair, and upkeep^7-9^. However, during AVS development, VIC-to-myofibroblast transition persists, leading to excessive ECM synthesis and remodeling, and consequent valve stiffening^8-10,22-27^. These pathological alterations of the aortic valve are recognized to be primary promoters of AVS development and progression^22-27^. Previous work by our lab and others showed that Trpv4 plays regulatory roles in myofibroblast activation and fibrosis development in other organs, suggesting that Trpv4 may act as a stiffness sensor in promoting AVS development^14,16-18,20,21,31^. Here, we focused on understanding whether Trpv4-mediated mechanotransduction specifically regulates VIC-myofibroblast activation as a function of matrix stiffness. Our current study sheds light on the Trpv4-dependent mechanism underlying VIC-myofibroblast activation through several major findings: (a) primary mouse VICs possess functional Trpv4 channels; (b) genetic ablation of Trpv4 blocked both soluble and mechanical factor-induced VIC-myofibroblast activation; (c) differential requirement of N-terminal Trpv4 domains for cellular force generation and VIC-myofibroblast activation; and (d) Trpv4 regulates matrix stiffness and TGFβ1-induced PI3K-AKT activity to modulate VIC-myofibroblast activation and cellular force generation.

During AVS development, VICs are likely subjected to a complex array of mechanical, physical, and soluble biochemical cues *in vivo* due to their interactions with other cell types, surrounding tissue matrix, and changing stiffness of the aortic valve matrix, which can modify their activation status^22-27,55,56^. Herein, we show that primary mouse VICs possess functional Trpv4 channels that can be modulated by its specific agonist and antagonist. Notably, stimulation of VICs by TGFβ1, a major factor linked to myofibroblast activation and fibrosis development^50-52^, augmented Trpv4 channel activity. Previous reports have shown increased Trpv4 expression and activity levels in fibrotic cells and tissue samples from numerous fibrotic conditions^14,20,40,41,44^. Moreover, it has been shown that porcine VICs grown under myofibroblast promoting medium conditions express increased Trpv4 proteins^41^. In this report, we further presented that Trpv4 deficiency suppressed both matrix stiffness and TGFβ1-induced cytoskeletal remodeling, as well as VIC-myofibroblast activation, which were restored by Trpv4 reintroduction. Notably, pretreatment of VICs by TGFβ1 further augmented matrix stiffness-induced VIC-myofibroblast activation. This may be due to increased activity of Trpv4 in VICs observed under TGFβ1 stimulation. Altogether, these results suggest that Trpv4 is directly involved in VIC-myofibroblast activation and also agree with a recent report showing the role of Trpv4 in regulating porcine VIC-myofibroblast activation and proliferation^41^. Using deletion strategies, we have now extended these findings. Here, we showed that N-terminal residues 30-130 in Trpv4 are indispensable for their profibrotic activity as revealed by its reliance on these residues for VIC-myofibroblast activation. In contrast, truncation of N-terminal residues 1-30 had no noticeable negative effect on VIC-myofibroblast activation. Since Trpv4 is sensitive to either surrounding matrix stiffness or TGFβ1 stimulation^20,23,26,27,41^, our finding that stiffer matrix or combination of stiffness plus TGFβ1 leads to increased VIC-myofibroblast activation in a Trpv4-dependent manner, further suggests a feed-forward mechanism in which a reciprocal functional interaction between Trpv4 and matrix stiffness propels AVS development and progression.

Cells, including fibroblasts, use adhesion molecules like integrin and ion channels like Trpv4 to sense changes in substrate stiffness associated with processes such as stress fiber development and cytoskeletal remodeling^47^. In our effort to identify a mechanosensing receptor and associated mechanisms by which mechanical signals are propagated into cells to drive cell differentiation, we previously reported that Trpv4 is involved in epithelial-to-mesenchymal transition, as well as lung and skin fibroblast differentiation^14,16-18,20,40^. Cytoskeletal remodeling is central to myofibroblast differentiation^14,16-18,20,40^. Notably, the generation of cellular force is primarily achieved by F-actin production, a key aspect of cytoskeletal remodeling^46,47^. Using traction force microscopy analysis, we showed that lack of Trpv4 function abrogated TGFβ1-induced force generation in VICs, which was re-established by Trpv4 reintroduction. These results suggest that Trpv4 is required for force generation in VICs under fibrogenic conditions. Furthermore, we presented that N-terminal regions 30-130 in Trpv4 are critical for its activity, as revealed by its reliance on these residues to generate cellular force. In contrast, deletion of residues 1-30 had no noticeable negative effect on force generation. Activation of the Trpv4 channel has been shown to require phosphatidylinositol 4,5-bisphosphate (PIP2) binding in the N-terminal region (residues 121-125) of Trpv4^45^. Deletion of N-terminal regions 30-130 was thought to interfere with PIP2-Trpv4 interaction, subsequently causing inhibition of activation of the channel. Whether similar PIP2-Trpv4 binding is also required for VIC-myofibroblast activation remains to be determined.

Herein, based on our gain-of and loss-of-function study, we found that Trpv4 is directly involved in regulating both matrix stiffness- and TGFβ1-induced phosphorylation of AKT, and generation of cellular force in VICs. These results suggest a role of Trpv4-PI3K axis in cellular force generation under stiff environment. Previously we reported that Trpv4 deletion inhibited both the matrix stiffness- and TGFβ1-induced activation of the PI3K-AKT pathway in skin epithelial cells^16,17^. The YAP/TAZ signaling pathway is a critical regulator of cell matrix mechanotransduction and signaling^58-62^. Previous reports from our lab and others linked Trpv4 to YAP/TAZ signaling in different cell types^16,17,41^. Currently, it is not well understood how calcium signaling through Trpv4 modulates YAP nuclear localization, and a more robust and precise pathway analysis is warranted. For example, efforts have been made to connect profibrotic pathways, such as PI3K and MAPK, and both the PI3K and the MAPK pathway have been recognized as bonafide targets of Trpv4^16,17,63^. Furthermore, it has been demonstrated that PI3K signaling modulates YAP/TAZ signaling in tumor cells, suggesting the potential for a Trpv4-PI3K-YAP/TAZ pathway in regulating VIC-myofibroblast activation^64^. However, with the currently available data, it remains speculative whether these pathways operate in parallel or are mutually exclusive. This underscores the necessity to more comprehensively identify the downstream targets of Trpv4 that influence VIC-myofibroblast activation. In this study, we used mouse VICs to explore the role of Trpv4 mechanotransduction in VICs. Therefore, to corroborate the proposed mechanisms, further experiments using human cells are warranted.

Studies investigating the function of Trpv4 in the aortic valve are scant. Although it has been found that aortic valve leaflets harvested from patients having aortic valve disease contain augmented levels of Trpv4 protein compared to that from patients with healthy hearts, not much is known about the mechanisms at play^57^. Recent evidence indicates that increased Trpv4 levels in the myocardium of diabetic rats mediate cardiac fibrosis^65^. Although aortic valve leaflet tissue was not examined, a similar process may be occurring in the context of AVS progression, as numerous studies from our lab and others point to ECM regulation of Trpv4 in fibroblasts^14,20,40,41,66^. Recent research has shown that Trpv4 expression is elevated in cardiomyocytes from aged mice compared to young mice and has linked Trpv4 to cardiomyocyte damage and possible participation in myofibroblast generation and fibrosis^66^.

In summary, the results of our study support the notion that a feed-forward functional interaction between Trpv4 and matrix stiffness leads to cytoskeletal remodeling and cellular force generation to regulate VIC-myofibroblast activation under AVS. As such, our study provides insights into the VIC-matrix interaction and Trpv4 mechanotransduction, which might be exploited in the development of targeted therapeutics for AVS.

## ACKNOWLEDGEMENTS

This work was supported by an NIH (R01EB024556) grant to Shaik O. Rahaman.

## AUTHOR CONTRIBUTIONS

SOR conceived the study, designed, and performed the experiments, analyzed data, and written and edited the MS. PM wrote the materials and method section of the manuscript and performed the experiments. SGR wrote and edited the manuscript. MM and BD maintained the animal colony, and KRS performed immunoblot experiments. All authors reviewed the manuscript and approved the final content of the manuscript.

## DATA AVAILABILITY

All data generated or analyzed during this study are included in this article.

## REFERENCES

1. Otto CM, Prendergast B. Aortic-Valve Stenosis-From Patients at Risk to Severe Valve Obstruction. N Engl J Med. 2014;371(8):744–756.

2. Yutzey KE, Demer LL, Body SC, Huggins GS, Towler DA, Giachelli CM, Hofmann-Bowman MA, Mortlock DP, Rogers MB, Sadeghi MM, Aikawa E. Calcific aortic valve disease: A consensus summary from the alliance of investigators on calcific aortic valve disease. Arterioscler Thromb Vasc Biol. 2014;34(11):2387–2393.

3. Schlotter F, Halu A, Goto S, Blaser MC, Body SC, Lee LH, Higashi H, DeLaughter DM, Hutcheson JD, Vyas P, Pham T, Rogers MA, Sharma A, Seidman CE, Loscalzo J, Seidman JG, Aikawa M, Singh SA, Aikawa E. Spatiotemporal Multi-Omics Mapping Generates a Molecular Atlas of the Aortic Valve and Reveals Networks Driving Disease. Circulation. 2018;138(4):377–393.

4. Goody PR, Hosen MR, Christmann D, Niepmann ST, Zietzer A, Adam M, Bönner F, Zimmer S, Nickenig G, Jansen F. Aortic valve stenosis: From basic mechanisms to novel therapeutic targets. Arterioscler Thromb Vasc Biol. 2020;(April):885–900.

5. Osnabrugge RL, Mylotte D, Head SJ, Van Mieghem NM, Nkomo VT, LeReun CM, Bogers AJ, Piazza N, Kappetein AP. Aortic stenosis in the elderly: disease prevalence and number of candidates for transcatheter aortic valve replacement: a meta-analysis and modeling study. J Am Coll Cardiol. 2013;62(11):1002–12.

6. Liang F, Wang X, Wang Q, Yan P, Yao L. Predilatation and paravalvular leakage risk in TAVR. Lancet. 2020;396(10251):600.

7. Liu AC, Joag VR, and Gotlieb AI (2007) The emerging role of valve interstitial cell phenotypes in regulating heart valve pathobiology. Am. J. Pathol 171, 1407–1418

8. Kendall RT and Feghali-Bostwick CA (2014) Fibroblasts in fibrosis: Novel roles and mediators. Front. Pharmacol 5 MAY, 1–13

9. Humphrey JD, Dufresne ER, and Schwartz MA (2014) Mechanotransduction and extracellular matrix homeostasis. Nat. Rev. Mol. Cell Biol 15, 802–812

10. Hinz B (2016) Myofibroblasts. Exp. Eye Res 142, 56–70

11. Kumar S. Cellular mechanotransduction: stiffness does matter. Nat Mater. 2014;13(10):918–920.

12. Tschumperlin DJ. Fibroblasts and the ground they walk on. Physiology (Bethesda). 2013;28(6):380–390.

13. Irianto J, Pfeifer CR, Xia Y, Discher DE. SnapShot: Mechanosensing Matrix. Cell. 2016;165(7):1820–1820.

14. Rahaman SO, Grove LM, Paruchuri S, Southern BD, Abraham S, Niese KA, Scheraga RG, Ghosh S, Thodeti CK, Zhang DX, Moran MM, Schilling WP, Tschumperlin DJ, Olman MA. TRPV4 mediates myofibroblast differentiation and pulmonary fibrosis in mice. J Clin Invest. 2014;124(12):5225–5238.

15. Matthews BD, Thodeti CK, Tytell JD, Mammoto A, Overby DR, Ingber DE. Ultra-rapid activation of TRPV4 ion channels by mechanical forces applied to cell surface beta1 integrins. Integr Biol (Camb). 2010;2(9):435–442.

16. Sharma S, Goswami R, Zhang DX, Rahaman SO. TRPV4 regulates matrix stiffness and TGFβ1-induced epithelial-mesenchymal transition. J Cell Mol Med. 2019;23(2):761–774.

17. Sharma S, Goswami R, Rahaman SO. The TRPV4-TAZ mechanotransduction signaling axis in matrix stiffness- and TGFβ1-induced epithelial-mesenchymal transition. Cell Mol Bioeng. 2019;12:139–152.

18. Sharma S, Goswami R, Merth M, Cohen J, Lei KY, Zhang DX, Rahaman SO. TRPV4 ion channel is a novel regulator of dermal myofibroblast differentiation. Am J Physiol Cell Physiol. 2017;312(5):C562–C572.

19. Bonnans C, Chou J, Werb Z. Remodelling the extracellular matrix in development and disease. Nat Rev Mol Cell Biol. 2014;15(12):786–801.

20. Goswami R, Arya RK, Sharma S, Dutta B, Stamov DR, Zhu X, Rahaman SO. Mechanosensing by TRPV4 mediates stiffness-induced foreign body response and giant cell formation. Sci Signaling. 2021, Nov 2: 14 (707): eabd4077.

21. Dutta B, Goswami R, Rahaman SO. TRPV4 Plays a Role in Matrix Stiffness-Induced Macrophage Polarization. Front. Immunol. 2020 Dec14;11:570195.

22. Go AS, Mozaffarian D, Roger VL, Benjamin EJ, Berry JD, Borden WB, et al. Heart disease and stroke statistics--2013 update: a report from the American Heart Association. American Heart Association Statistics Committee and Stroke Statistics Subcommittee. Circulation. 2013;127(1):e6–e245.

23. Ma H, Killaars AR, DelRio FW, Yang C, Anseth KS. Myofibroblastic activation of valvular interstitial cells is modulated by spatial variations in matrix elasticity and its organization. Biomaterials. 2017;131:131–144.

24. Quinlan AMT, Billiar KL. Investigating the role of substrate stiffness in the persistence of valvular interstitial cell activation. J Biomed Mater Res A. 2012;100(9):2474–82.

25. Bruschi G, Maloberti A, Sormani P, Colombo G, Nava S, Vallerio P., Casadei F, Bruno J, Moreo A, Merlanti B, Russo CD, Oliva F, Klugmann S, Giannattasio C. Arterial Stiffness in Aortic Stenosis: Relationship with Severity and Echocardiographic Procedures Response. High Blood Press Cardiovasc Prev. 2017;24(1):19–27.

26. Determinants and clinical significance of aortic stiffness in patients with moderate or severe aortic stenosis. Saeed S, Saeed N, Grigoryan K, Chowienczyk P, Chambers JB, Rajani R. Int J Cardiol. 2020;315:99–104.

27. Redirecting valvular myofibroblasts into dormant fibroblasts through light-mediated reduction in substrate modulus. Wang H, Haeger SM, Kloxin AM, Leinwand LA, Anseth KS. PLoS One. 2012;7(7):e39969.

28. Dutta B, Arya RK, Goswami R, Alharbi MO, Sharma S, Rahaman SO. Role of macrophage TRPV4 in inflammation. Lab Invest. 2020;100(2):178–185.

29. Blakney AK, Swartzlander MD, Bryant SJ. The effects of substrate stiffness on the in vitro activation of macrophages and in vivo host response to poly(ethylene glycol)-based hydrogels. J Biomed Mater Res A. 2012;100(6):1375–1386.

30. Hind LE, Dembo M, Hammer DA. Macrophage motility is driven by frontal-towing with a force magnitude dependent on substrate stiffness. Integr Biol (Camb). 2015;7(4):447–453.

31. Arya RK, Goswami R, Rahaman SO. Mechanotransduction via a TRPV4-Rac1 signaling axis plays a role in multinucleated giant cell formation. J Biol Chem. 2020 Dec1;296:100129.

32. Garcia-Elias A, Mrkonjić S, Jung C, Pardo-Pastor C, Vicente R, Valverde MA. The TRPV4 channel. Handb Exp Pharmacol. 2014;222:293–319.

33. Everaerts W, Nilius B, Owsianik G. The vanilloid transient receptor potential channel TRPV4: from structure to disease. Prog Biophys Mol Biol. 2010;103(1):2–17.

34. Liedtke W. Molecular mechanisms of TRPV4-mediated neural signaling. Ann N Y Acad Sci. 2008;1144:42–52.

35. Thorneloe KS, Cheung M, Bao W, Alsaid H, Lenhard S, Jian MY, Costell M, Maniscalco-Hauk K, Krawiec JA, Olzinski A, Gordon E, Lozinskaya I, Elefante L, Qin P, Matasic DS, James C, Tunstead J, Donovan B, Kallal L, Waszkiewicz A, Vaidya K, Davenport EA, Larkin J, Burgert M, Casillas LN, Marquis RW, Ye G, Eidam HS, Goodman KB, Toomey JR, Roethke TJ, Jucker BM, Schnackenberg CG, Townsley MI, Lepore JJ, Willette RN. An orally active TRPV4 channel blocker prevents and resolves pulmonary edema induced by heart failure. Sci Transl Med. 2012;4(159):159ra148.

36. Everaerts W, Zhen X, Ghosh D, Vriens J, Gevaert T, Gilbert JP, Hayward NJ, McNamara CR, Xue F, Moran MM, Strassmaier T, Uykal E, Owsianik G, Vennekens R, De Ridder D, Nilius B, Fanger CM, Voets T. Inhibition of the cation channel TRPV4 improves bladder function in mice and rats with cyclophosphamide-induced cystitis. Proc Natl Acad Sci U S A. 2010;107(44):19084–9.

37. Masuyama R, Vriens J, Voets T, Karashima Y, Owsianik G, Vennekens R, Lieben L, Torrekens S, Moermans K, Vanden Bosch A, Bouillon R, Nilius B, Carmeliet G. TRPV4-Mediated Calcium Influx Regulates Terminal Differentiation of Osteoclasts. Cell Metab. 2008;8:257–265.

38. Al-Shammari H, Latif N, Sarathchandra P, McCormack A, Rog-Zielinska EA, Raja S, Kohl P, Yacoub MH, Peyronnet R, Chester AH. Expression and function of mechanosensitive ion channels in human valve interstitial cells. PLoS One. 2020;15(10 October):1–18. E0240532.

39. Jia X, Xiao C, Sheng D, Yang M, Cheng Q, Wu J, Zhang S. TRPV4 Mediates Cardiac Fibrosis via the TGF-β1/Smad3 Signaling Pathway in Diabetic Rats. Cardiovasc Toxicol. 2020;20(5):492–499.

40. Goswami, R., Cohen, J., Sharma, S., Zhang, D. X., Lafyatis, R., Bhawan, J., and Rahaman, SO. TRPV4 ion channel is associated with scleroderma. J. Invest Dermatol. 2016;137(4):962–965.

41. Batan D, Peters DK, Schroeder ME, Aguado BA, Young MW, Weiss RM, Anseth KS. Hydrogel cultures reveal Transient Receptor Potential Vanilloid 4 regulation of myofibroblast activation and proliferation in valvular interstitial cells. FASEB J. 2022 ;36(5):e22306.

42. Bouchareb, R., Lebeche, D. Isolation of Mouse Interstitial Valve Cells to Study the Calcification of the Aortic Valve In Vitro. J. Vis. Exp. (171), e62419, doi:10.3791/62419 (2021).

43. Oh RS, Haak AJ, Smith KMJ, Ligresti G, Choi KM, Xie T, Wang S, Walters PR, Thompson MA, Freeman MR, Manlove LJ, Chu VM, Feghali-Bostwick C, Roden AC, Schymeinsky J, Pabelick CM, Prakash YS, Vassallo R, Tschumperlin DJ. RNAi screening identifies a mechanosensitive ROCK-JAK2-STAT3 network central to myofibroblast activation. J Cell Sci. 2018;131(10):jcs209932.

44. Adapala RK, Katari V, Teegala LR, Thodeti S, Paruchuri S, Thodeti CK. TRPV4 Mechanotransduction in Fibrosis. Cells. 2021;10(11):3053.

45. Garcia-Elias A, Mrkonjic S, Pardo-Pastor C, Inada H, Hellmich UA, Rubio-Moscardó F, Plata C, Gaudet R, Vicente R, Valverde MA. Phosphatidylinositol-4,5-biphosphate-dependent rearrangement of TRPV4 cytosolic tails enables channel activation by physiological stimuli. Proc Natl Acad Sci U S A. 2013;110(23):9553–8.

46. Defife KM, Jenney CR, Colton E, Anderson JM, Disruption of filamentous actin inhibits human macrophage fusion. FASEB J. 1999;13, 823–832.

47. Gavara N, Chadwick RS, Relationship between cell stiffness and stress fiber amount, assessed by simultaneous atomic force microscopy and live-cell fluorescence imaging. Biomech. Model. Mechanobiol. 2016;15, 511–523.

48. Larue L, Bellacosa A. Epithelial-mesenchymal transition in development and cancer: role of phosphatidylinositol 3’ kinase/AKT pathways. Oncogene. 2005; 24: 7443–7454.

49. Tan WJ, Tan QY, Wang T, Lian M, Zhang L, Cheng ZS. Calpain 1 regulates TGF-β1-induced epithelial-mesenchymal transition in human lung epithelial cells via PI3K/Akt signaling pathway. Am J Transl Res. 2017; 9: 1402–1409.

50. Biernacka A, Dobaczewski M, Frangogiannis NG. TGF-β signaling in fibrosis. Growth Factors. 2011; 29: 196–202.

51. Akhurst RJ, Hata A. Targeting the TGFβ signalling pathway in disease. Nat Rev Drug Discov. 2012; 11: 790–811.

52. Wynn TA, Ramalingam TR. Mechanisms of fibrosis: therapeutic translation for fibrotic disease. Nat Med. 2012; 18: 1028–1040.

53. Rubashkin MG, Cassereau L, Bainer R, DuFort CC, Yui Y, Ou G, Paszek MJ, Davidson MW, Chen YY, Weaver VM. Force engages vinculin and promotes tumor progression by enhancing PI3K activation of phosphatidylinositol (3,4,5)-triphosphate. Cancer Res. 2014; 74: 4597–4611.

54. Leight JL, Wozniak MA, Chen S, Lynch ML, Chen CS. Matrix rigidity regulates a switch between TGF-β1-induced apoptosis and epithelial-mesenchymal transition. MolBiol Cell. 2012; 23: 781–791.

55. Rabkin E, Aikawa M, Stone JR, Fukumoto Y, Libby P, Schoen FJ. Activated interstitial myofibroblasts express catabolic enzymes and mediate matrix remodeling in myxomatous heart valves. Circulation. 2001 Nov 20;104(21):2525–32.

56. Rabkin-Aikawa E, Farber M, Aikawa M, Schoen FJ. Dynamic and reversible changes of interstitial cell phenotype during remodeling of cardiac valves. J Heart Valve Dis. 2004;13(5):841–7.

57. Al-Shammari H, Latif N, Sarathchandra P, McCormack A, Rog-Zielinska EA, Raja S, Kohl P, Yacoub MH, Peyronnet R, Chester AH. Expression and function of mechanosensitive ion channels in human valve interstitial cells. PLos One. 2000;15,1–8.

58. Low BC, Pan CQ, Shivashankar GV, Bershadsky A, Sudol M, Sheetz M. YAP/TAZ as mechanosensors and mechanotransducers in regulating organ size and tumor growth. FEBS Letters. 2014; 588: 2663–2670.

59. Aragona M, Panciera T, Manfrin A, Giulitti S, Michielin F, Elvassore N, Dupont S, Piccolo S. A mechanical checkpoint controls multicellular growth through YAP/TAZ regulation by actin-processing factors. Cell. 2013; 154: 1047–1059.

60. Dupont S, Morsut L, Aragona M, Enzo E, Giulitti S, Cordenonsi M, Zanconato F, Le Digabel J, Forcato M, Bicciato S, Elvassore N, Piccolo S. Role of YAP/TAZ in mechanotransduction. Nature. 2011; 474: 179–183.

61. Piersma B, Bank RA, Boersema M. Signaling in Fibrosis: TGF-β, WNT, and YAP/TAZ Converge. Front Med (Lausanne). 2015; 2: 59.

62. Liu F, Lagares D, Choi KM, Stopfer L, Marinkovic A, Vrbanac V, Probst CK, Hiemer SE, Sisson TH, Horowitz JC, Rosas IO, Fredenburgh LE, Feghali-Bostwick C, et al. Mechanosignaling through YAP and TAZ drives fibroblast activation and fibrosis. American journal of physiology. Lung cellular and molecular physiology. 2015; 308: L344–357.

63. Grove LM, Mohan ML, Abraham S, Scheraga RG, Southern BD, Crish JF, Naga Prasad SV, Olman MA (2019) Translocation of TRPV4-PI3Kγ complexes to the plasma membrane drives myofibroblast transdifferentiation. Sci. Signal 12, 1–15.

64. Zhao Y, Montminy T, Azad T, Lightbody E, Hao Y, SenGupta S, Asselin E, Nicol C, and Yang X (2018) PI3K positively regulates YAP and TAZ in mammary tumorigenesis through multiple signaling pathways. Mol. Cancer Res 16, 1046–1058.

65. Jia X, Xiao C, Sheng D, Yang M, Cheng Q, Wu J, Zhang S. TRPV4 Mediates Cardiac Fibrosis via the TGF-β1/Smad3 Signaling Pathway in Diabetic Rats. Cardiovasc Toxicol. 2020;20(5):492–499.

66. Jones JL, Peana D, Veteto AB, Lambert MD, Nourian Z, Karasseva NG, Hill MA, Lindman BR, Baines CP, Krenz M, Domeier TL. TRPV4 increases cardiomyocyte calcium cycling and contractility yet contributes to damage in the aged heart following hypoosmotic stress. Cardiovasc Res. 2019;115(1):46–56.

